# Auditory perceptual history is propagated through alpha oscillations

**DOI:** 10.1101/553222

**Authors:** Hao Tam Ho, David C. Burr, David Alais, Maria Concetta Morrone

## Abstract

Perception is a proactive, *predictive* process, where the brain uses accumulated experience to make best guesses about the world to test against sensory data, updating the guesses as new experience is acquired. Using novel behavioral methods, the present study demonstrates the role of alpha rhythms in communicating past perceptual experience. Participants were required to discriminate the ear of origin of brief sinusoidal tones that were presented monaurally at random times within uncorrelated dichotic white noise masks. Responses were not constant over time but oscillated rhythmically towards one or the other ear, at ~9 Hz. Importantly, these alpha oscillations occurred only for trials preceded by a target tone to the same ear, either on the previous trial or two trials ago. Tones to the other ear elicited no oscillations. These results suggest that the phase-resetting of oscillations (necessary to measure them by behavioral means) is contingent on the congruency on the previous stimuli. Overall the results strongly implicate behavioral oscillations with the processes of serial dependence and hence with memory formation.

## Introduction

It has been long known that perception depends heavily on expectations and perceptual experience. Helmholtz [1] introduced the concept of *unconscious inference*, suggesting that perception is at least partly *inferential*, or *generative*, and Gregory [2] described perception as a series of *hypotheses* to be verified against sensory data, using many compelling illusions to support this notion. On this view perception is a proactive, *predictive* process, where the brain uses accumulated experience to make best guesses about the world to test against sensory data, updating the guesses as new experience is acquired.

Recent studies using *serial dependence* demonstrate clearly the action of predictive perception and provide a means of quantitative study: under many conditions, the appearance of images in a sequence depends strongly on the stimulus presented just prior to the current one. Judgments of orientation [3], numerosity [4], motion [5], facial identity or gender [6, 7], beauty and even perceived body size [8] are strongly biased towards the previous image. Serial biases are also observed in audition for pitch discrimination [9, 10]. Sequential effects can last up to minutes [11], showing that perception does not rely solely on instantaneous stimulation, but also on predictions, or *priors*, conditioned by events over a long time-course. Storage of prior information necessarily implicates memory processes.

The neuronal mechanisms underlying serial dependence are largely unknown. It is assumed that the predictions are generated at mid-high levels of analysis, and fed back to early sensory areas, whose activity in turn feeds forward to add to the accumulated knowledge and shape future predictions [12–16]. Little is known, however, about how this information is propagated, or the nature of the underlying neural mechanisms. One possibility is that recursive propagation and updating of stored prior experience is related to low-frequency neural oscillations [17–19]. We have recently shown, in both audition [20] and vision [21], that *sensitivity* (accuracy) and *criterion* (response bias) are not constant but oscillate rhythmically over time, at different frequencies: theta for sensitivity and alpha for criterion, suggesting separate mechanisms [20, 21]. The alpha oscillations in criterion are consistent with an increasing number of EEG findings showing an association between criterion shifts and modulations of alpha power and phase [18, 22–25].

Oscillations in bias could plausibly reflect the action of predictive mechanisms, possibly via reverberation of recursive error propagation within a generative framework. VanRullen and Macdonald [26] have proposed an oscillatory mechanism by which past perceptual history may be stored in short-term memory as a reverberatory “perceptual echo”. The reverberation should affect the predictive mechanism that biases perceptual decisions, giving rise to sequential effects. Given the oscillatory nature of the reverberation, behavioral oscillations in criteria, observed in different domains and tasks [20, 21, 27], may be modulated or even gated by the history of the previous presented stimuli. We test this idea by measuring behavioral oscillations in criterion in an auditory discrimination task, analyzing the series based on the congruency of previous auditory stimuli. The results demonstrate that perceptual oscillations occur only for stimuli that are congruent with previous stimuli.

## Results

### Stimulus history affects response bias but not sensitivity

Participants were required to identify in two-alternative forced-choice the ear of origin of a brief monaural near-threshold tone embedded within a 2-second burst of dichotic white noise (Fig. 1A). We first looked for serial dependence effects in sensitivity and response bias, as measured respectively by d-prime (*d’*) and decision criterion *(c*) (Eq. 1&2). Stimulus history had no influence on observer sensitivity (*p* = 0.07 and *p* = 0.56 for 1- and 2-back, respectively), as may be expected as target location was totally independent of the previous trial. However, criterion was affected by stimulus history (Fig. 1B). Although there was no significant 1-back effect (FDR adjusted *p* = 0.86), there was a strong 2-back effect of response biases towards the previously presented stimulus (*p* = 0.0002). The effects remained significant for stimuli three and four trials back (*p* = 0.008 and 0.009 respectively) but were not significant five trials back (*p* = 0.85).

**Figure 1.**
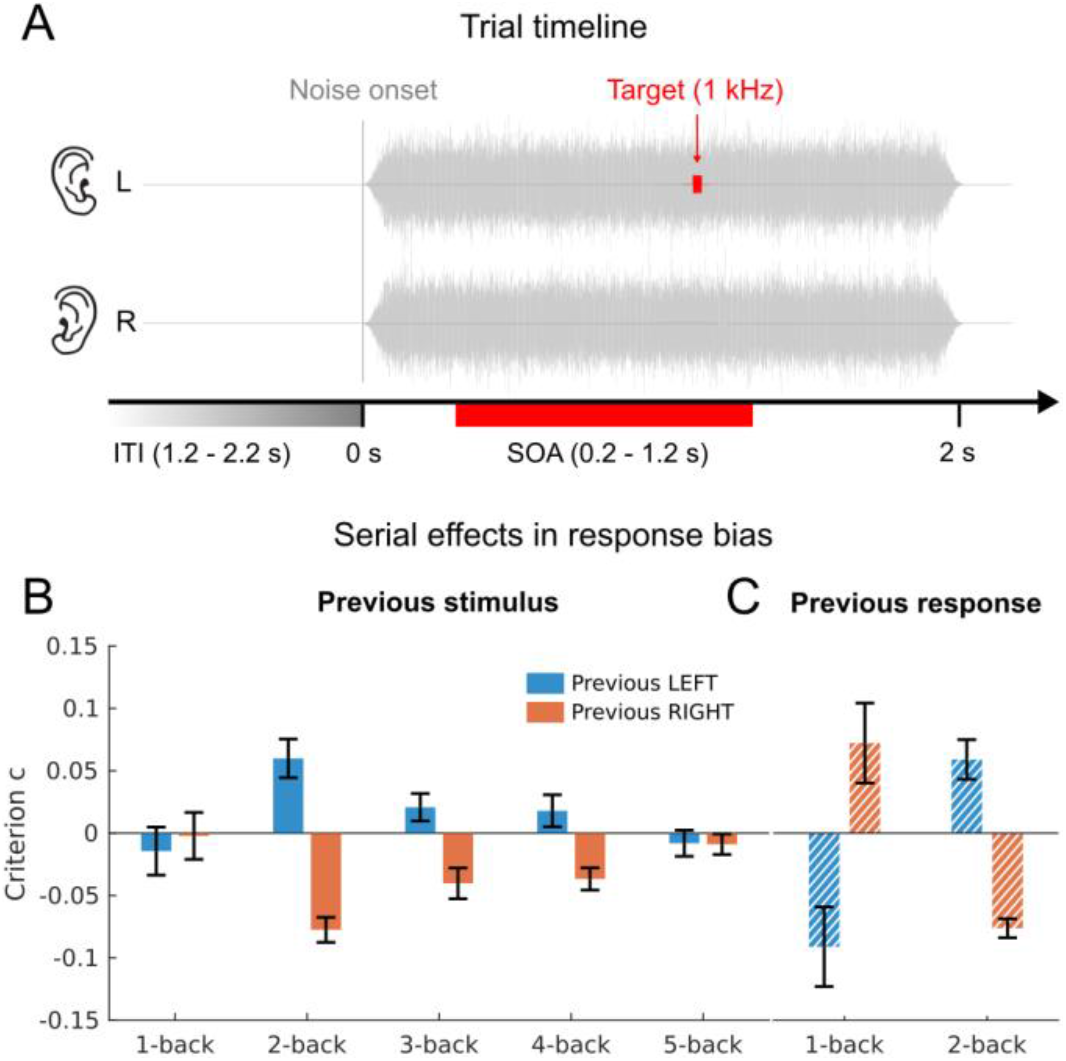
**(A)** Schematics of a trial. On each trial, white noise was presented simultaneously to both ears for 2 s. A pure tone of 1 kHz and 10 ms duration was delivered with equiprobability to the left or right ear, at a SOA randomly selected from an interval of 0.2-1.2 s post noise onset. The inter-trial interval varied randomly between 1.2-2.2 s. **(B)** Results of the overall serial-dependence analysis. Group mean response bias (as measured by the decision criterion *c*, Eq. 2) contingent on the ear of origin of the preceding 1 to 5 stimuli. The difference between the contingent left and right (blue and red bars, respectively) are significant for the 2-, 3- and 4-back stimuli (FDR corrected). **(C)** Group mean response bias contingent on the response 1-2 trials back. The difference between the contingent left and right (blue and red bars, respectively) is significant for both 1- and 2-back (FDR corrected). Sensitivity (*d’*, not plotted) showed no significant sequential effects for either 1- or 2-back trials.

Analysis of the dependency of the *previous response* (Fig. 1C) revealed a *negative* influence of one trial back, marginally significant after FDR correction (*p* = 0.053). Although not statistically robust, this repulsive effect could result from response switching [28], and explain the lack of stimulus-based serial dependence in 1-back trials. Response dependence on two trials back (which should not be affected by response switching) showed the same assimilative aftereffect found for the stimulus-based analysis (*p* = 0.0001, FDR corrected). Trials further in the past (not shown) had no significant effect (all *p’s* > 0.05).

### Oscillation of response bias but not sensitivity

Figures 2 A&B show the variation over time in sensitivity (Fig. 2A) and criterion (Fig. 2B), computed by binning aggregate data as a function of target SOA from noise onset. It is evident that criterion oscillates strongly and regularly and can be well fit by a pure sinusoidal function with frequency of 9.4 Hz, shown by the grey curve in Fig. 2B. However, sensitivity does not show a rhythmic periodicity and no sinusoidal function fit well the temporal series (Fig. 2C). To evaluate the goodness of fit of each sinusoidal function, we compared the explained variance (*R*^*2*^) of the real data for every frequency from 4-12 Hz in steps of 0.1 Hz with the *R*^*2*^ of the best fit of the surrogate shuffled data at any frequency within the range (Figs. 2C-E: see methods). For sensitivity (orange line), no frequency produced a fit near the 95%-confidence threshold (dotted line). However, the criterion data (green line) showed significant modulations between 9.2 and 9.6 Hz, with a strong peak at 9.4 Hz (*R*^*2*^ = 0.15). The phase of the 9.4 Hz oscillation at trial onset relative to the noise burst onset was 179° ± 16° SEM (by bootstrap). For sensitivity (left), the explained variance is clearly not significant, while for criterion (right) the best fit is higher than 99.1% of the surrogate best fits, giving a sign-test significance of *p* = 0.009.

**Figure 2.**
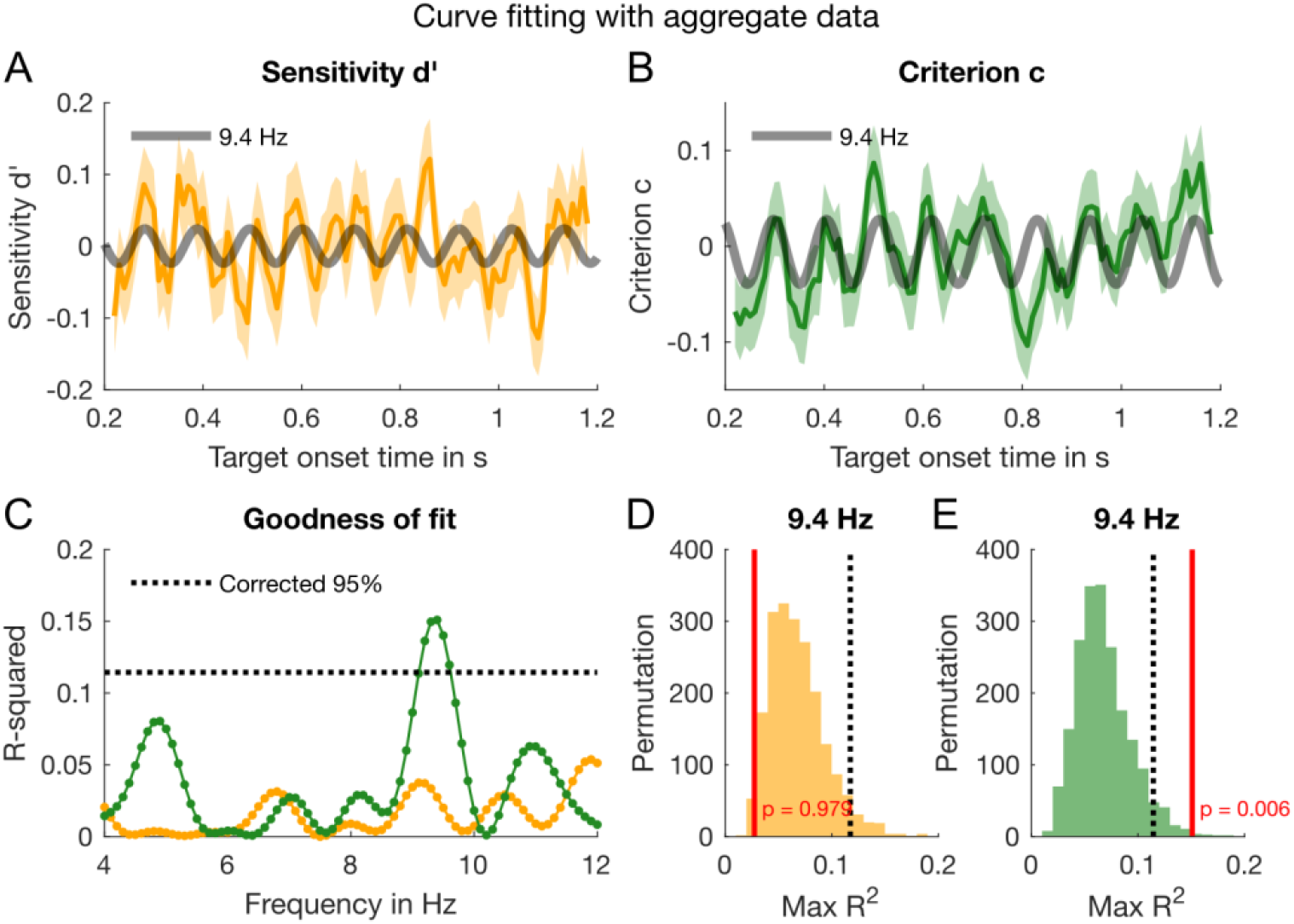
Results of the curve fitting analysis. **(A)** The yellow line shows the time course of the detrended sensitivity (*d’*) based on the aggregate data (pooling all trials across the 14 participants and binning with a rectangular smoothing window of 20 ms that shifted in 10 ms steps). The data are smoothed the data for display all statistical analyses used non-smoothed data. The shaded yellow and green areas enveloping the lines represent bootstrapped ±1 SEM. The grey curve depicts a 9.4 Hz oscillation fitted to the sensitivity data. **(B)** The analysis of the criterion (*c*) data (green line) followed the same binning and curve fitting procedure. **(C)** The goodness of fit (*R*^*2*^) for all sinusoids from 4-12 Hz, in steps of 0.1 Hz. The dotted black line represents the 95 percentiles of the permutation distribution depicted in E, which we take as the threshold for significance. *R*^*2*^ for sensitivity (yellow line) never reaches significance, while that for criterion does for the range of 9.1 – 9.6 Hz, highest at 9.4 Hz. **(D-E)** Illustration of the permutation method: we shuffled the aggregate data 2,000 times, fitted the shuffled data with the best frequency over the range, and calculated the distributions of *R*^*2*^ for sensitivity (D) and criterion (E) distributions. The red lines show the *R*^*2*^ of the fit to the original data at 9.4 Hz. The *p*-values of the sign test is given by the proportion of permuted *R*^*2*^ greater than the original *R*^*2*^ (red lines).

The results shown in Figure 2 were based on an aggregate data analysis, by pooling all trials across subjects, and fitting sinusoids over the entire duration. In a complementary analysis, we analyzed the data of individual participants (without data binning) and evaluated their coherence as a group using the regression analysis in Eq. 5, applying a GLM to the *accuracy* (correct or incorrect) and *response bias* (‘left’ or ‘right’) of single trials. The results corroborate those of the aggregate data analysis. Figure 3A plots the amplitude spectrum for oscillations in *accuracy* (an approximation of sensitivity), and Figure 3B that for *response bias* (an approximation of criterion). Response bias (Fig. 3B) shows a strong peak around 9.4 Hz, reinforcing the curve-fitting results for criterion in Figure 2C. A similar permutation procedure as for the aggregate analysis yielded the corrected *p*-values plotted in Figures 3D&E. For response bias, the oscillation at 9.4 Hz is significant (*p* = 0.027) after correction for multiple comparisons with maximal statistics, illustrated in Figure 3F. While there are several peaks in the amplitude spectrum for accuracy (Fig. 3A), none was significant after multiple comparison correction (Fig. 3D).

**Figure 3.**
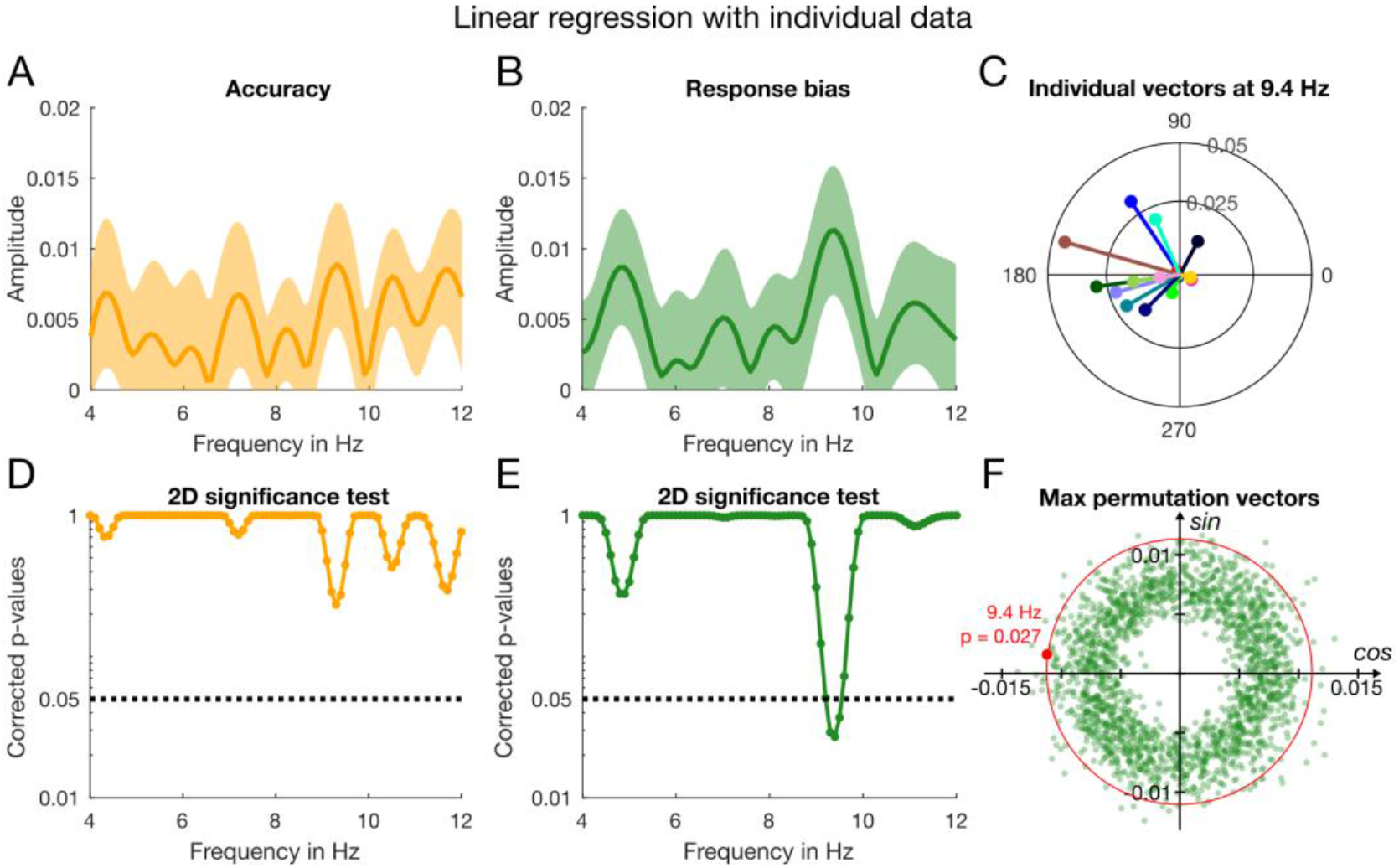
Results of the linear regression analysis based on individual data without binning. **(A)** The yellow line represents the amplitude spectrum for accuracy computed from the vectorial average of the GLM estimates of *β*_1_ and *β*_2_ across participants. The shaded area around the line indicates ±1 SEM. **(B)** Amplitude spectrum of response bias based on the same analyses as for accuracy. **(C)** Individual 2D vectors (*β*_1_, *β*_2_) at 9.4 Hz for response bias. The length and direction of the line indicate the amplitude and phase (referred to time of noise onset). **(D)** The results of the 2D permutation test for accuracy for the frequency range of interest, 4-12 Hz. We corrected for multiple comparisons using maximal statistics illustrated in F. **(E)** Corrected *p*-values for response bias obtained by the same 2D permutation as for accuracy. **(F)** Illustration of the 2D permutation test by which the *p*-values in D&E were computed. The sign test is based on the proportion of maximal permutation vectors that exceed the group mean (outside the red circle passing through the group mean, shown by the red dot).

Figure 3C shows the individual vectors for response bias at 9.4 Hz, with vector angle showing the phases of individual participants at noise onset. The vectors are tightly clustered around a phase angle of 172° ± 16° SEM, similar to the phase angle we obtained from the curve fitting analysis with the aggregate data (179°).

### Oscillations in response bias are driven by stimulus history

Having established the existence of rhythmic fluctuations in criterion in both the aggregate and individual data, we investigated the dependence of the oscillations on the previous stimulus, using the same two analysis techniques. We separated the trials into two groups, based on whether the previous stimulus had been presented to the same ear (congruent) or different ear (incongruent), and analyzed for oscillations in sensitivity and criterion. The sensitivity data showed no significant oscillations at any frequency, either for congruent or incongruent data (see Figure S1 in supplemental material). Perhaps this was to be expected as there were no oscillations in the total data set.

Figure 4 shows the results of the curve-fitting analysis of the aggregate data for criterion. Congruent trials (Fig. 4A, dark green line), showed a good fit at 9.4 Hz (thick grey line, *R*^*2*^ = 0.15, *p* = 0.01), while for incongruent trials the goodness of fit did not approach significance at any frequency ((Fig. 4B, light green line). Figure 4C shows that only for congruent trials did the goodness of fit survive the multiple comparison correction, and only for frequencies between 9.1 and 9.6 Hz (*p* = 0.01 at 9.4 Hz). The phase of this 9.4-Hz oscillation for congruent trials was 180° ± 17° (SEM evaluated by bootstrap). For incongruent trials, the phase at 9.4 Hz was 179° ± 34°.

**Figure 4.**
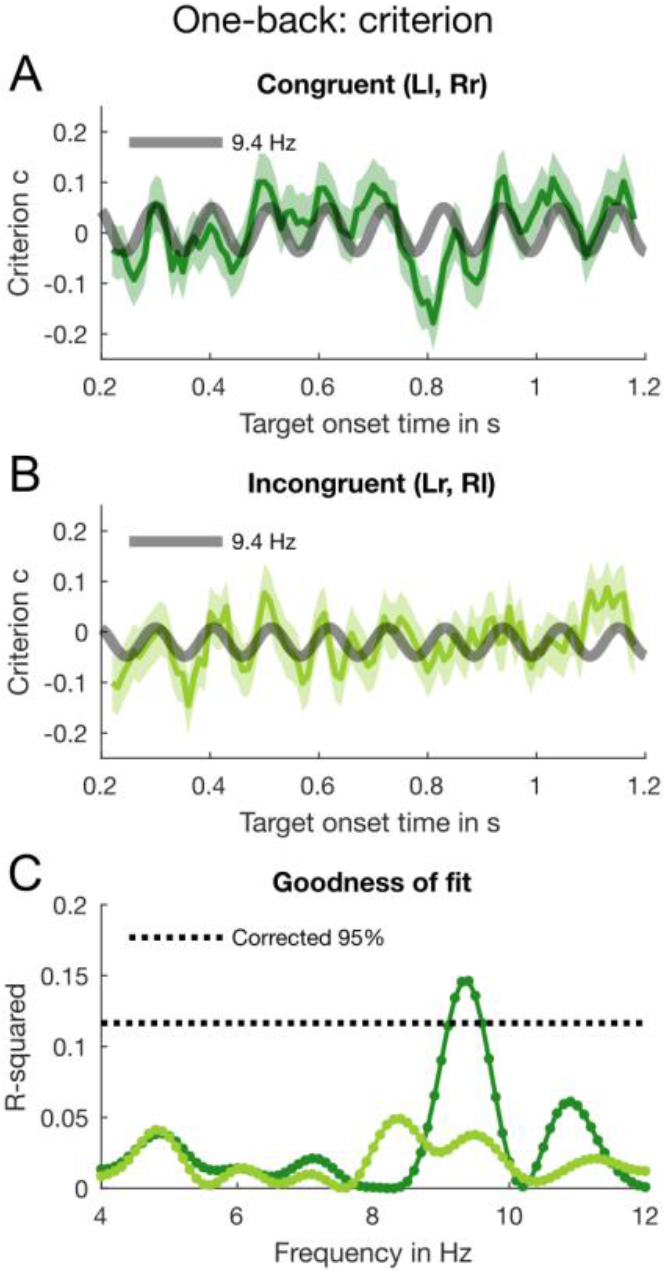
Results of the one-back analysis for criterion with aggregate data. **(A)** The dark green line shows the binned *congruent* trials (data smoothed for display purposes only). The error bars indicate ±1 SEM obtained by bootstrapping the aggregate data 2,000 times. The thick grey line represents the 9.4-Hz oscillation, which we fitted to the criterion data. **(B)** The *incongruent* trials submitted to the same binning, curve fitting and bootstrapping procedure as congruent trials. **(C)** The goodness of fit for congruent (dark green line) and incongruent trials (light green line) at all tested frequencies from 4-12 Hz in 0.1-Hz steps. The black dotted line indicates the 95-percentile of the distribution of maximal *R*^*2*^ obtained by permuting the individual trials.

Again, we examined the individual subject data using the regression analysis, separately for congruent and incongruent trials. As for the aggregate data, the congruent trials (dark green line) yielded a large peak around 9.4 Hz with *A* = 0.03 ± 0.009 (Fig. 5A). At this frequency, the amplitude for the incongruent trials is much reduced, with *A* = 0.02 ± 0.01 (Fig. 5B). Inspection of the vector plots shows a tight cluster around a mean phase angle (at noise onset) of 164° ± 13° SEM for congruent trials (Fig. 5C), but a greater dispersion for incongruent trials (mean phase 177° ± 18° SEM: Fig. 5D). The results of the 2D permutation test plotted in Figure 5E show that the only frequencies to survive the strict multiple comparison correction were around 9.4 Hz (dark green line, congruent trials) with *p*= 0.045. In contrast, incongruent trials showed no significant frequencies (light green line).

**Figure 5.**
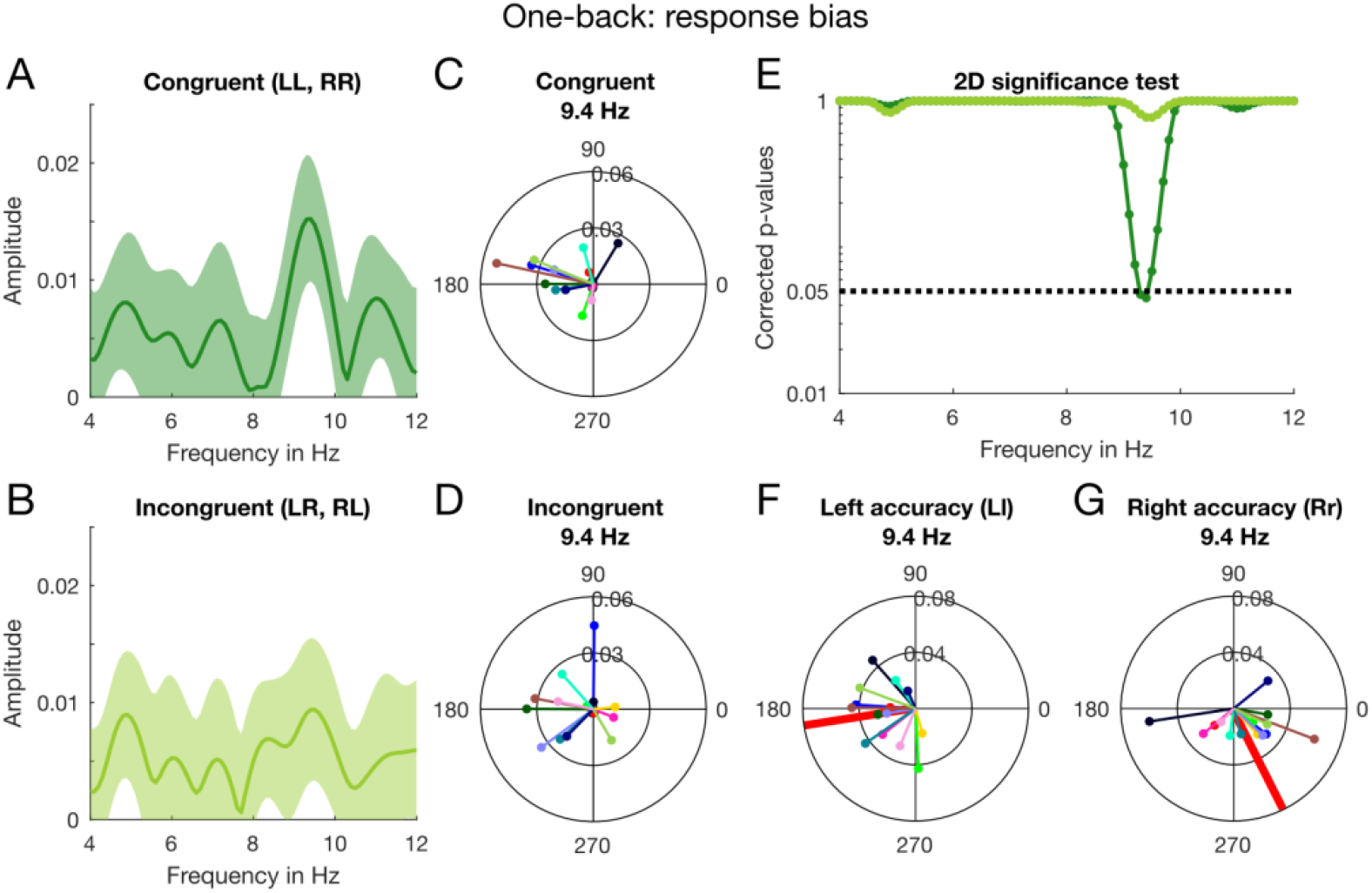
Results of the one-back analysis for response bias with individual subject data. **(A)** Amplitude spectrum for the congruent trials computed from the individual estimates of *β*_1_ and *β*_2_ averaged across participants. The shaded area around the dark green line indicates ±1 SEM. **(B)** By the same method, we computed the amplitude spectrum with incongruent trials. **(C)** Individual phase and amplitude vectors (at noise onset) based on congruent trials at 9.4 Hz. **(D)** Individual vectors for incongruent trials at 9.4 Hz. **(E)** The 2D significance test (see Fig. 3F) was done for every frequency between 4-12 Hz in 0.1-Hz steps. The dark and light green lines depict the corrected *p*-values for congruent and incongruent trials, respectively. The black dotted line indicates *α* = 0.05 (corrected for multiple comparisons). **(F)** As a post-hoc test, we split the congruent trials further into trials that contained a left or right target and tested their phase relationship using circular statistics. At 9.4 Hz observers showed very strong phase coherence for both ears. Here, we plot the individual phase and amplitude vectors for congruent trials containing a target in the left ear. The thick red line indicates the direction of the mean vector (with unit length) which is close to 180°. **(G)** The individual phase and amplitude vectors at 9.4 Hz for congruent trials containing a right target. The direction of the mean vector is ~300°.

As our previous work suggested that sensitivity may oscillate out of phase in the two ears [20], we performed a post-hoc regression analysis of the accuracy of congruent trials separately for ear of origin. Figure 5F&G show significant phase consistencies for *both* ears for the *congruent* trials (when analyzed separately) at 9.4 Hz. Using the Watson-Williams test (circular analogue to a two-sample *t*-test; [29]), we confirmed that the group phase distributions for left- and right-ear accuracy in the congruent trials were significantly different (*p* < 0.05, Bonferroni corrected) between 9.1 and 9.7 Hz, peaking at around 9.4 Hz. Figure 6G shows the individual vectors at 9.4 Hz for congruent trials: left targets have a mean direction 189° ± 12°, right targets 304° ± 14° (Fig. 5G). The mean difference is 115°, broadly consistent with an antiphase relationship. Thus, the lack of oscillations in the total dataset (Figs. 2&3) may reflect cancellation of out-of-phase left and right ear oscillations, consistent with our previous research [20].

**Figure 6.**
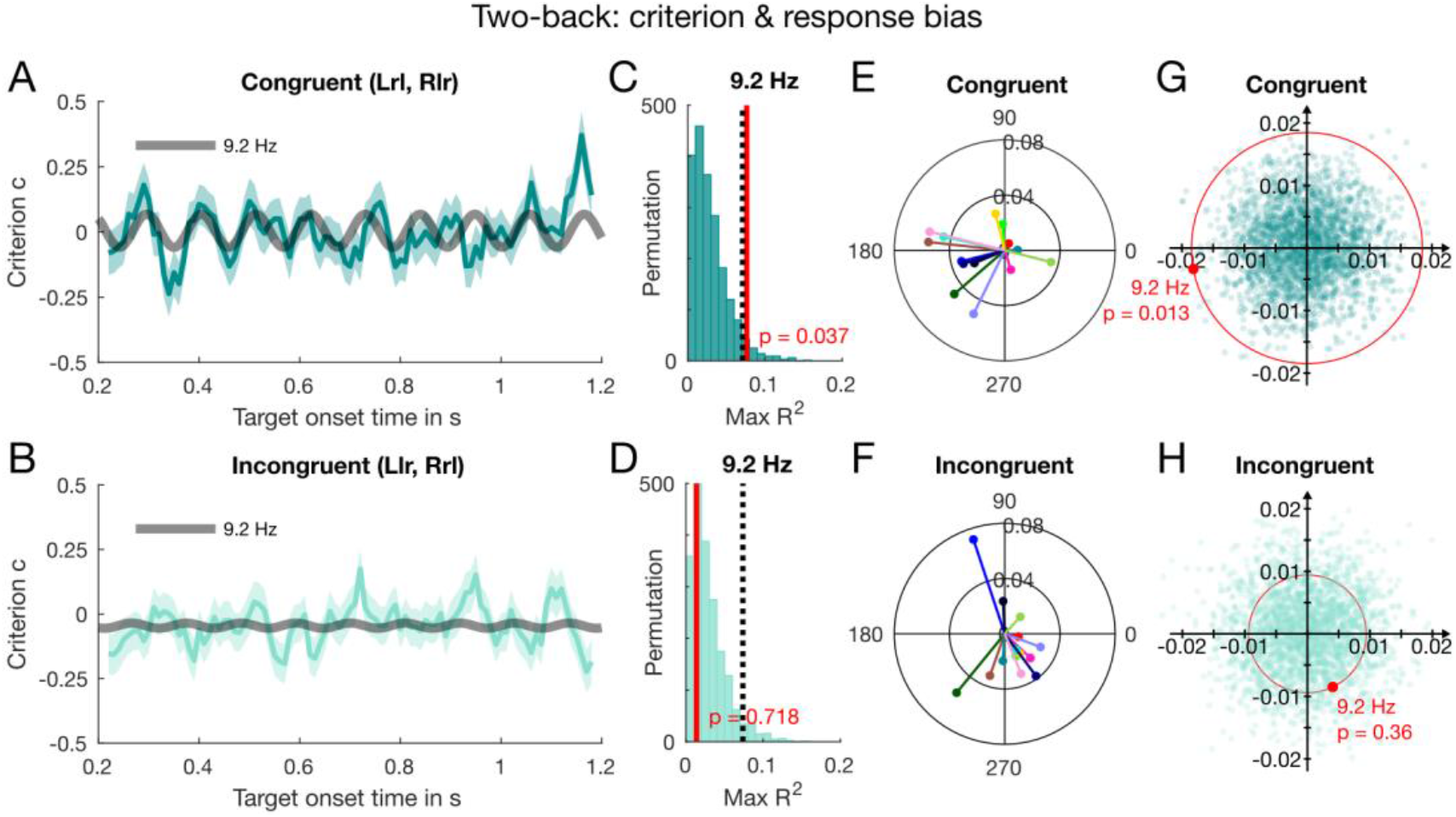
Results of the two-back analysis for criterion with aggregate data of trials *incongruent* with the previous trial, either congruent or incongruent with stimuli 2 trials back. **(A)** The dark cyan line shows the binned *congruent* trials (error bars indicate ±1 SEM), with the dark grey thick line showing the best fitting sinusoid (9.2 Hz, *R*^*2*^ = 0.08) over the range of 9.1 to 9.6 Hz. **(B)** The binned incongruent data fitted with the same frequency, 9.2 Hz. **(C)** The *R*^*2*^ obtained at 9.2 Hz (thick red line) was compared against the goodness-of-fit of the surrogate shuffled data (dark cyan histogram), binned and fitted as the original data. **(D)** The same permutation test for the incongruent condition was not significant. **(E)** The individual vectors in the congruent condition at 9.2 Hz, with subjects color-coded as in Figures 4&6. Their phases cluster around a similar phase as in the congruent 1-back (Fig. 5C). **(F)** The incongruent phases at 9.2 Hz have mean phase that bears no relation to either the congruent or incongruent mean phase in the 1-back (Figs. 5C&D). **(G)** The result of the 2D permutation test for the congruent condition is significant, consistent with the result of the aggregate data analysis shown in (C). **(H)** The 2D permutation result for the incongruent condition is not significant, also consistent with the aggregate result in (D).

An obvious question is whether the oscillations in criteria depend on the current stimulus being the same as the previous stimulus or the previous response. This is difficult to test, as responses were 75% correct, and therefore strongly correlated with stimuli. However, we ran the curve-fitting analysis for current stimuli contingent on the previous response. On this analysis, the main peak remained at 9.4 Hz, but was not significant (see Fig. S2 in supplemental material). This suggests – but does not prove – that the trials needed to be coherent with previous stimuli rather than responses. No other analyses, including coherent and incoherent error trials (where responses are different from stimuli) yielded significant results (see Fig. S3 in supplemental material). However, these error analyses necessarily result in greatly reduced numbers of trials (about 25%), making significance difficult to reach.

### Duration of the serial effect

As the serial dependence analysis showed strong 2-back effects (Fig. 2B), we tested whether measurable oscillations near 9.4 Hz were contingent on stimuli presented two trials back. We examined further the 1-back incongruent data, which showed no significant oscillations (Fig. 5), dividing it into trials congruent or incongruent with stimuli presented two trials back. As we expected oscillations to be weak and near the frequency found for 1-back trials, we confined our analysis window to a limited region of 9.1-9.6 Hz (for both the original and shuffled data). Figure 6A shows a significant oscillation in the 2-back congruent data, best fitted by a sinusoid of ~9.2 Hz (*R*^*2*^ = 0.08, *p* = 0.037). There was no modulation at this frequency in the 2-back incongruent data (*R*^*2*^ = 0.02, *p* = 0.7, Fig. 6B).

We examined the individual group coherence at 9.2 Hz with the same regression analysis as before (Eq. 5), for 2-back congruent (Fig. 6E) and 2-back incongruent (Fig. 6F) trials. The phases of the congruent trials cluster around 190° ± 18°, similar to that of the 1- back congruent data (Fig. 5C), while that for the incongruent trials is 295° ± 15°, which bears no relation to the mean phase of either the 1-back congruent or incongruent condition (Figs. 5C&D). Figures 6G&H show the results of the 2D permutation test at 9.2 Hz, which are consistent with the results of the aggregate data analysis. Very few points (dark cyan dots in Fig. 6G) from the permutation distribution (*p* = 0.013) exceed the group mean vector (thick red dot) in the congruent condition, compared with the incongruent condition, *p* = 0.36 (Fig. 6H). Taken together, the results suggest that the 9.2-Hz oscillation lasts at least two trials.

## Discussion

To build and maintain a stable and coherent percept in a noisy and ambiguous world, observers take advantage of past information to anticipate forthcoming sensory input. While there is a good deal of behavioral evidence in favor of this predictive account of perception, little is known about the underlying neural mechanisms. The current study suggests that predictive perception may be implemented rhythmically through alpha-band oscillations. Identifying the ear of origin of a weak tone was strongly biased by previous stimuli, two or even three trials before the current trial. Although the immediately past trials had no average serial effect, we observed a strong rhythmic fluctuation in bias (or *criterion*) at ~9.4 Hz that was critically dependent on stimulus history: oscillations occurred *only* when a stimulus had been presented to the same ear as the current one, either one or two trials previously.

The strong dependency of oscillations on stimulus coherency suggests that they play a fundamental role in the propagation of predictive information. That oscillations occurred only for stimuli presented to the same ear as the previous stimuli suggests that stimuli somehow initiate or gate oscillatory processes that last until the next trial, and even the one after. These oscillations may be related to the “perceptual echo” suggested by VanRullen and Macdonald [26].

It is far from clear how the oscillations last for so long, up to two trials, ~6-8 seconds. It is of course possible that the stimulus itself sets in train an oscillation that is phase-reset by the onset of the next trial’s noise burst, possibly with re-amplification. Alternatively, it may be that the stimulus does not actually start an oscillation, but somehow sensitizes or primes the circuitry to oscillate, so that when next noise burst of the next trial arrives the oscillations begin. A third possibility is that oscillations are always present, and a stimulus to that ear primes their phase-resetting by the next noise burst: without a phase-reset, the oscillations will not be synchronous with noise onset, and therefore will not be measurable by our behavioral technique. At this stage it is impossible to distinguish between the three alternatives: however, the results clearly implicate alpha oscillations in the propagation of information about perceptual history.

The behavioral oscillations probably result from rhythmical variation of neuronal sensitivity or response gain. But why should oscillations in gain affect criterion rather than sensitivity? Both this and our previous [20] study suggest that the oscillations may have opposite starting phase in each ear. As sensitivity (*d’*) can be calculated only after combination of hits and false alarms of both ears, the out-of-phase modulations should cancel each other out. Indeed, post-hoc tests of congruent trials separated for ear of origin of the signal (Figs. 5F&G), together with evidence from our previous study [20], supports the suggestions that monaural oscillations in sensitivity are in antiphase in each ear. On the other hand, combining the counter-phased modulation for *criterion* (the sum of hits and false alarms, oppositely signed for each ear) will not annul the modulations, but cause them to add together in the binaural measurements. On this interpretation modulation of *criterion* is associated with perceptual changes [30, 31], rather than with modulation of a decision boundary. Overall, our interpretation is consistent with the notion of modulation of neuronal response gain induced by phase-resetting endogenous neuronal rhythms, which results in criterion modulation in our experimental design.

We have described the stimulus contingency in terms of ear of origin. However, it is important to note that stimuli confined to one ear will be perceived as originating from that side of space. It is therefore possible that the interaction between consecutive stimuli may not occur at the peripheral monaural level, but at later representation of space. In other words, the oscillation could be between spatial regions, rather than between ears. This explanation would be consistent with several studies in vision reporting behavioral oscillations between spatial locations [32] and between spatial objects [33]. Although the modulations in visual sensitivity were in the theta band, they show that cyclic alternation of spatial sensitivity could be a general perceptual strategy. Further evidence along these lines comes from Lozano-Soldevilla and VanRullen’s [34] demonstration of opposite EEG phase in the two hemispheres of “perceptual echoes”, 10 Hz reverberations in response to visual stimulation. It seems reasonable to expect similar effects in audition.

Why did our experiments reveal no average positive serial dependence on the immediately previous trial (1-back), as is normally observed in studies of serial dependence? There are (at least) two possible, non-mutually exclusive explanations. One is that forced-choice paradigms can lead to sequential response biases, such as alternation [28], which would tend to cancel out positive serial dependence based on the previous stimulus, but have no average effect for trials two-back. The response-based analysis showed a strong negative serial dependency, consistent with response alternation, potentially cancelling perceptual positive serial dependence. Another possible reason is that the stimuli may have caused both positive serial dependence and negative adaptation aftereffects. Negative aftereffects tend to be shorter lived than assimilative dependencies, and may therefore cancel out only 1-back, not 2-back trials [11]. Whatever the reason for the lack of positive serial dependence in the averaged 1-back results, our study shows that oscillations may be a more sensitive signature of memory-based perceptual effects than simply looking at average results. Many competing effects could reduce or annul average serial dependence effects, without affecting rhythmic, time-dependent oscillations.

In summary, we showed that when discriminating the ear of origin of a brief sinusoidal target within dichotic white noise, decision criteria oscillate at an alpha rhythm, but only when the previous target had been presented to the same ear. To account for these findings, we propose that presentation of a target potentiates ear-specific, alpha-rhythm reverberations, that bias perceptual decisions rhythmically, presumably by rhythmic modulation of response gain. The exact mechanisms of this process are far from clear, but the results show that alpha oscillations play a major role in propagating expectations and past perceptual history to affect current perception. It would be interesting to study these effects further with neurophysiological techniques, such as EEG, MEG or Functional Near Infrared Imaging.

## Materials and Methods

### Participants

Eighteen healthy adults with normal hearing took part in the experiment. Three were excluded for imbalanced left and right ear auditory thresholds and one for very long reaction times (2.5 standard deviations from the group mean). Of the remaining 14 participants (mean age 21.14 ± 4.22), 4 were male and 2 left-handed. All participants provided written, informed consent. The study was approved by the Human Research Ethics Committees of the University of Sydney. We based our sample size estimations on our previous study [20], which showed oscillations in auditory perceptual performance before, and other studies on similar behavioral rhythms in vision [32, 36], without running a formal power analysis.

### Experimental procedure

Participants sat in a dark room and listened to auditory stimuli via in-ear tube-phones (ER-2, Etymotic Research, Elk Grove, Illinois) with earmuffs (3M Peltor 30 dBA) to isolate external noise. On each trial, 2 seconds of binaural broadband white-noise (randomly generated each trial and uncorrelated between the two ears) were presented together with the monaural target tone. The target (1000 Hz, 10 ms) was delivered randomly with equal probability to either ear during the 2-s noise burst, within 0.2–1.2 s from noise onset. For each ear, the target intensity was kept near individual thresholds (75 % accuracy), using an accelerated stochastic approximation staircase procedure [37, 38]. Participants reported the *ear of origin* of the tone via button press (ResponsePixx, Vpixx Technologies, Saint-Bruno, Quebec). The next trial started after a silent inter-trial interval (ITI) of random duration between 1.2–2.2 s. Participants completed 2,800 trials (40 blocks of 70 trials with rests between blocks, and no feedback) after a practice block of 20 trials with feedback. Stimuli were presented using the software *PsychToolbox* [39] in conjunction with *DataPixx* (Vpixx Technologies) in MATLAB (Mathworks, Natick, Massachusetts). Trials were excluded if the response occurred before the target onset or after the noise offset, or if the reaction time (RT) exceeded the 99 % confidence interval of that individuals’ RT.

### Signal detection theory

To separate sensitivity and response bias, we computed d-prime, *d’*, and the decision criterion, *c*, using Eq. 1 and 2 from *signal detection theory* (SDT) [40, 41]. *d*′ is given by the difference between z-transformed hit and false alarm rates:

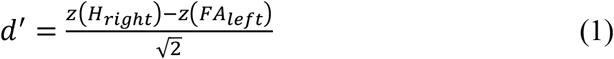

The bias of the responses was defined positive for the left ear:

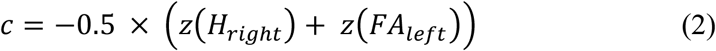

### Aggregate data analysis

The trials were pooled across all 14 participants and grouped into one hundred 10-ms bins, from 0.2 to 1.2 s post noise onset. The mean number of trials per bin was 151 ± 24 for the left-target condition and 152 ± 25 for the right-target condition. For each bin, we computed *d*’ and *c* as above and fitted the time-series with a sinusoidal function given by:

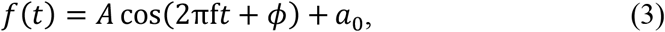

where *t* is time, *a_0_* a constant and *A* and ϕ amplitude and phase of the sinusoidal fit, all free parameters. The frequency parameter *f* was fixed between 4 and 12 Hz in 0.1 Hz steps and a non-linear least-squares method was used to obtain the best fit for each tested frequency (a standard implementation in MATLAB with 400 iterations in total). Sensitivity displayed a decreasing non-linear trend over time (see also [20]), which we removed before curve fitting. Detrending was not required for the criterion time series. The goodness of fit *R*^*2*^ was used to test the significance of every fit by applying a permutation procedure [42]: responses of each individual trial were shuffled over all SOAs to generate 2,000 surrogate datasets, which we submitted to the same binning and curve fitting procedure as the original data. To correct for multiple comparisons, we determined the maximal *R*^*2*^ for every surrogate dataset irrespective of frequency. This resulted in a distribution of 2,000 maximal *R*^*2*^ (Figs. 2D&E), against which we compared each fit to the original dataset. Any frequency that exceeded the 95 percentile of the maximal-*R*^*2*^ distribution (dotted line in Fig. 2C) was considered significant. We also estimated the variability in the original aggregate data by applying the bootstrap method. We randomly selected the same number of trials (with replacement) from the original data 2,000 times, and each surrogate dataset was submitted to the same binning and curve fitting procedure as above.

### Individual and group analyses

To examine the individual data for oscillations and evaluate their coherence across subjects, we used an approach based on single trials (for similar approaches, see [27, 43]). The response *y*_i_ (*i* = 1, 2… *n*, where *n* is the total number of trials) to a target at time *t*_i_ (i.e., the interval from noise onset to target onset in seconds) is modelled by the linear combination of harmonics at each tested (angular) frequency as follows:

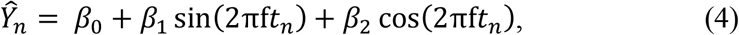

where 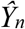 represents the predicted responses and *β*_0_, *β*_1_ and *β*_2_ are fixed-effect regression parameters estimated using the *linear* least-squares method of MATLAB (*fitlm* function from the *Statistics and Machine Learning* toolbox). This *general linear model* (GLM) model estimates the regression parameters adequately when the sampling rate is uniform across the time series. As this condition may not always be met at the individual subject level, we included a third independent regressor containing information about the stimulus:

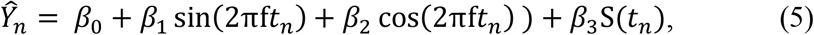

where *S* is the stimulus at time *t* and takes the value −1 or +1 for left and right target respectively. For sensitivity, *y*_i_ = 1 for correct and *y*_i_ = 0 for incorrect responses, and for response bias, *y*_i_ = 1 for a ‘right’ response and 0 for ‘left’. Although binary responses can be modelled with a logit or probit link function, non-linear transform of 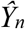 made no difference in our study (for further discussion, see [44]), so we performed no transformation of 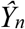.

The significance of the model fit in Eq. 5 was evaluated with a two-dimensional permutation test: we shuffled the SOAs of each individual’s trials to create 2,000 surrogate datasets per subject and fitted each dataset with the model described in Eq. 5. As with the original data, the resulting *β*_1_ and *β*_2_ were averaged across subjects for every frequency tested. This yielded a joined distribution of 2,000 surrogate means for each frequency from 4- 12 Hz in 0.1-Hz steps. To correct for *multiple comparison*, we determined the *maximal vector* of each joined distribution irrespective of the frequency. This resulted in a joint distribution of 2,000 maximal vectors, against which we compared the original group mean.

## Acknowledgements

The research was supported by the Australian Research Council Discovery Project (DP150101731) and the European Research Council (FPT/2007 – 2013) under grant agreements 338866 “Ecsplain” and 832813 “GenPercept”.

## Author Contributions

H.T.H. conducted the experiment; M.C.M., D.C.B., D.A. and H.T.H designed the experiment and wrote the paper.

## Supplemental information

**Figure S1.**
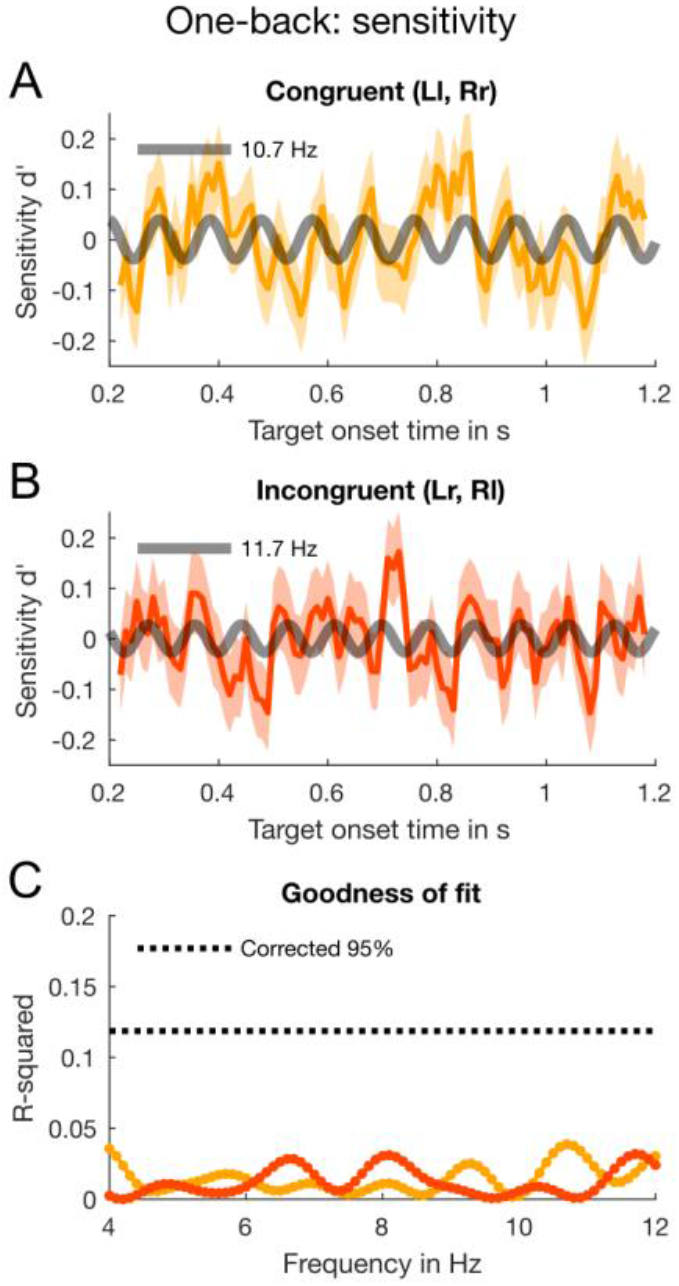
Results of the one-back analysis for sensitivity with aggregate data. **(A)** The yellow line shows the binned *congruent* trials (data smoothed for display purposes only). The error bars indicate ±1 SEM obtained by bootstrapping the aggregate data 2,000 times. The thick grey line shows the best-fitting sinusoidal function at 10.7-Hz oscillation. This fit was not significant. **(B)** The *incongruent* trials submitted to the same binning, curve fitting and bootstrapping procedure as congruent trials. The best fit (grey line) was at 11.7 Hz. Again, this fit was not significant. **(C)** The goodness of fit for congruent (dark green line) and incongruent trials (light green line) at all tested frequencies from 4-12 Hz in 0.1-Hz steps. The black dotted line indicates the 95-percentile of the distribution of maximal *R*^*2*^ obtained by permuting the individual trials. None of the tested frequency approached the 95% confidence threshold.

**Figure S2.**
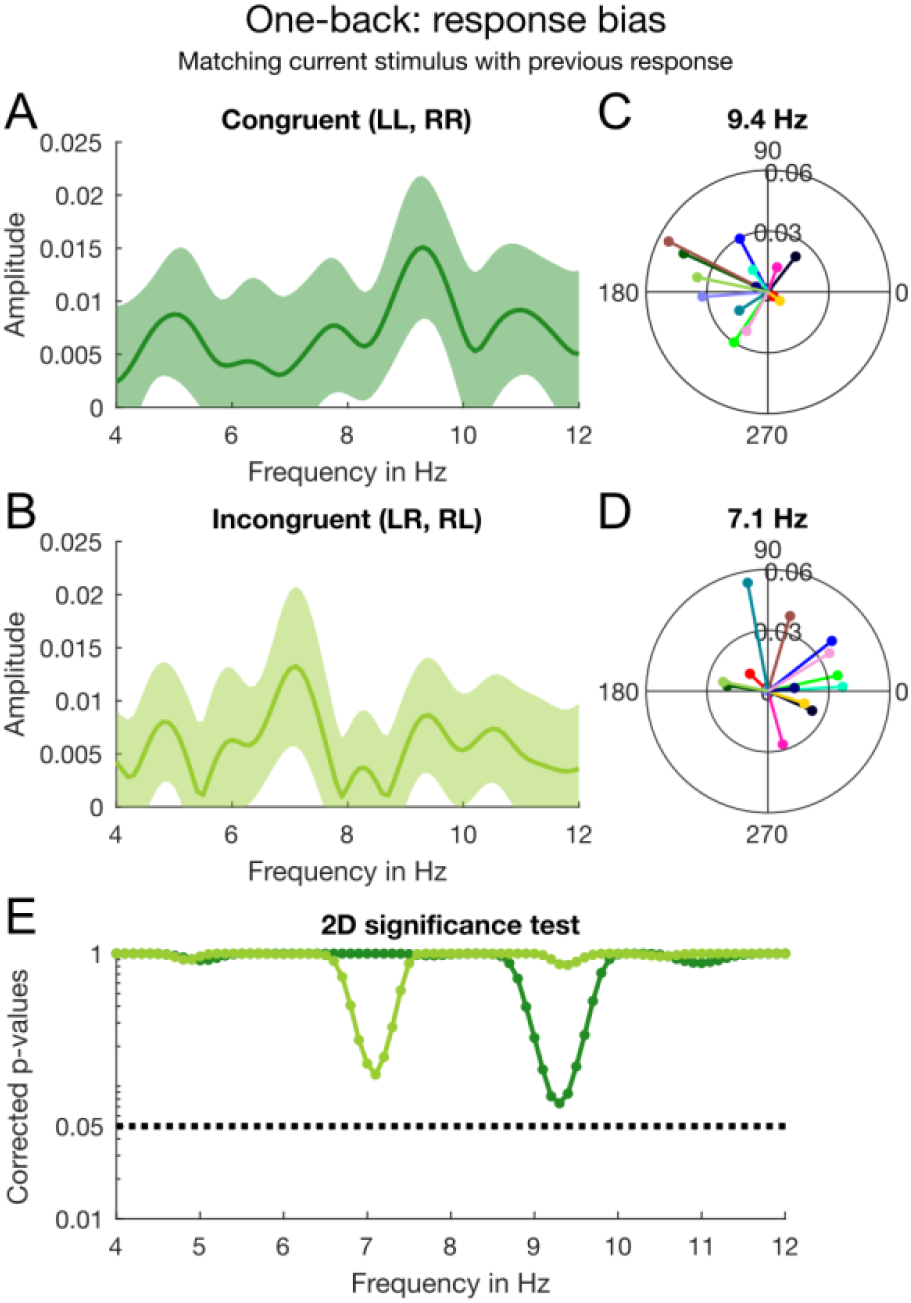
Results of the one-back analysis for response bias with individual subject data contingent on whether the *current stimulus* matches the *previous response*. **(A)** Amplitude spectrum for the congruent trials (current stimulus matches previous response) computed from the individual estimates of *β*_1_ and *β*_2_ averaged across participants. The shaded area around the dark green line indicates ±1 SEM. **(B)** By the same method, we computed the amplitude spectrum with incongruent trials (current stimulus *does not match* previous response). **(C)** Individual phase and amplitude vectors (at noise onset) based on congruent trials at 9.4 Hz. **(D)** Individual vectors for incongruent trials at 9.4 Hz. **(E)** The 2D significance test (see Fig. 3F) was done for every frequency between 4-12 Hz in 0.1-Hz steps. The dark and light green lines depict the corrected *p*-values for congruent and incongruent trials, respectively. The black dotted line indicates *α* = 0.05 (corrected for multiple comparisons).

**Figure S3.**
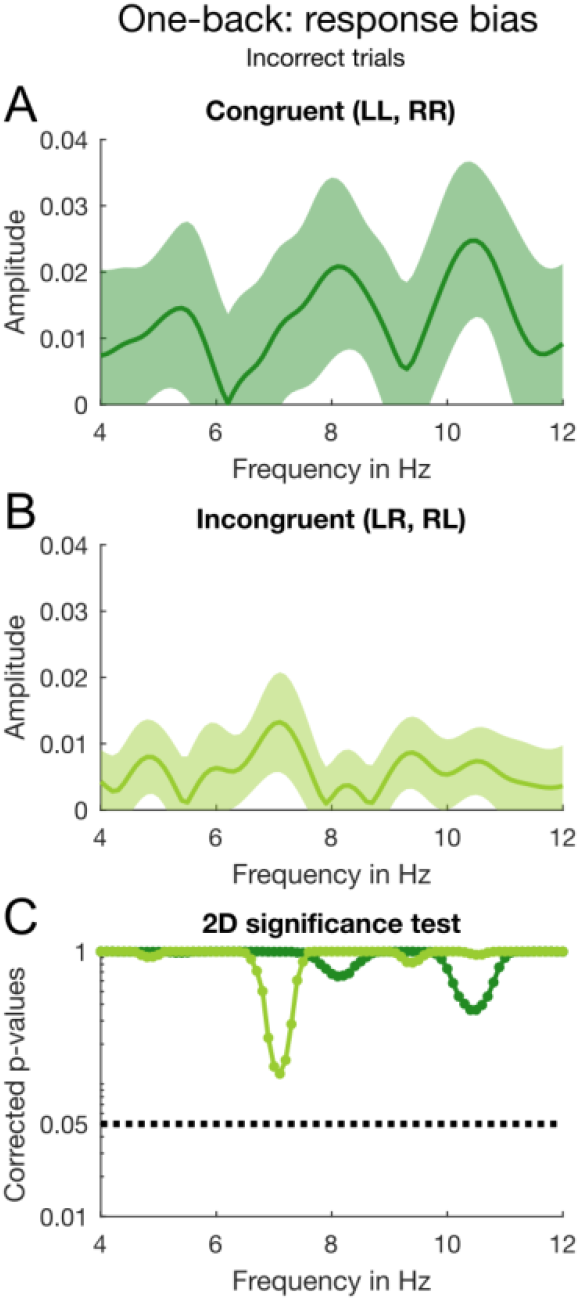
Results of the one-back analysis for response bias with individual subject data contingent on whether the *current stimulus* matches the *previous response*. This is a subset of the data shown in Fig. S2 containing only *incorrect* trials. **(A)** Amplitude spectrum for the congruent trials (current stimulus matches previous response) computed from the individual estimates of *β*_1_ and *β*_2_ averaged across participants. The shaded area around the dark green line indicates ±1 SEM. **(B)** By the same method, we computed the amplitude spectrum with incongruent trials (current stimulus *does not match* previous response). **(C)** Individual phase and amplitude vectors (at noise onset) based on congruent trials at 9.4 Hz. **(D)** Individual vectors for incongruent trials at 9.4 Hz. **(E)** The 2D significance test (see Fig. 3F) was done for every frequency between 4-12 Hz in 0.1-Hz steps. The dark and light green lines depict the corrected *p*-values for congruent and incongruent trials, respectively. The black dotted line indicates *α* = 0.05 (corrected for multiple comparisons).

